# A mouse model to study Río Negro virus pathogenesis: a neglected member of the Venezuelan equine encephalitis virus complex

**DOI:** 10.64898/2026.01.29.702336

**Authors:** Luciana A. Fassola, Yamila Gazzoni, Glenda D. Martin Molinero, Soledad De Olmos, María F. Triquell, Alicia Degano, Marianela C. Serradell, María E. Rivarola, Sergio R. Oms, Marta S. Contigiani, Adriana Gruppi, Guillermo Albrieu-Llinás

## Abstract

Río Negro virus (RNV) is an enzootic alphavirus and a member of the Venezuelan equine encephalitis virus (VEEV) complex. Despite its wide circulation in South America, RNV remains a neglected pathogen with no established wild-type animal model to study its pathogenesis. In this study, we developed a lethal mouse model using 18-day-old wild-type C57BL/6 mice to characterize the systemic and neurological features of RNV infection. Following subcutaneous inoculation, RNV exhibited rapid systemic dissemination with a brief and low-titer viremic phase and high viral loads in lymphoid tissues, pancreas, brains, and lungs. Notably, infected mice developed progressive neurological signs, including ataxia and hindlimb paralysis, culminating in 100% lethality. Histopathological analysis revealed significant damage, highlighted by a striking collapse of the splenic architecture, inflammatory and remodeling changes in the lungs, and prominent inflammatory infiltrates with neurodegenerative changes in the brain. The splenic disruption was further evaluated by immunofluorescence analysis of the spleen, which showed a consistent loss of compartmentalization, characterized by an atypical infiltration of CD8+ T cells into B-cell follicles. The terminal stage of disease was characterized by extensive neuroinflammation and neurodegeneration. Histological examination of the brain revealed meningoencephalitis, robust astrogliosis, and widespread somatodendritic and terminal degeneration, particularly clustered around blood vessels. These findings were supported by cytokine analysis of brain homogenates, which showed a significant upregulation of IFN-γ, IL-6, and MCP-1/CCL2 during symptomatic stages. Collectively, these findings establish a reproducible, non-genetically modified animal model that reveals the pathogenic potential of RNV in the context of immune immaturity characteristic of early life. By identifying these specific pathological and neuroinflammatory markers, our study provides a foundational experimental framework to investigate the mechanisms underlying RNV emergence and host-pathogen interactions within the VEEV complex.

**Author summary:** Many viruses circulate silently in nature, hidden within animal populations, until environmental or social changes bring them into contact with humans. The Río Negro virus is one such example. Despite being closely related to the better-known Venezuelan equine encephalitis virus and showing evidence of circulation in South America, RNV has remained largely overlooked by the scientific community and public health authorities. In this study, we established a new experimental model using infant mice that allowed us to observe how the virus spreads and causes damage. We found that the virus rapidly reaches the brain and other vital organs, causing severe inflammation in the brain, inflammatory changes in the lungs, and a breakdown of the immune system’s organization in the spleen. By using a genetically unmodified model, we were able to observe the infection in a host with an intact but naturally immature immune system. This approach avoids the artificial conditions of genetic engineering while providing a more realistic window into how host age and developmental stages can influence the outcome of neglected viral infections. We believe our work is a first step toward understanding how this overlooked virus emerges and highlights the need to monitor and prepare for “silent” pathogens that could pose a risk to public health in an era of intensifying human activity and ecological pressure.

## Introduction

The global resurgence of epidemics over recent decades has drawn renewed attention to arthropod-borne viruses (arboviruses). Nevertheless, many arboviruses remain poorly studied, particularly in developing regions, where they often circulate silently or cause mild disease and therefore fall outside global surveillance priorities. As a result, their eco-epidemiology and pathogenic potential remain incompletely understood.

Venezuelan equine encephalitis virus (VEEV) belongs to a complex of antigenically related alphaviruses that circulate throughout the Americas (https://talk.ictvonline.org/). Traditionally, VEEV variants have been classified as epidemic/epizootic or enzootic, based on their transmission cycles and association with outbreaks of disease in humans and equines. While epidemic variants are responsible for periodic outbreaks of severe disease, enzootic viruses have long been considered of limited pathogenic relevance, as they typically produce low viremia in equines and are not associated with large epidemics (1). However, human exposure to enzootic VEEV strains has been documented, including in urban settings, and infections may range from asymptomatic to mild febrile illness that is clinically indistinguishable from other arboviral infections, such as dengue. Consequently, the burden and pathogenic potential of enzootic VEEV infections are likely underestimated (2).

In recent years, increasing attention has been paid to enzootic arboviruses that, while not associated with large outbreaks, may cause significant disease under specific ecological or host-related conditions (3). Within this context, enzootic viruses of the Venezuelan equine encephalitis virus (VEEV) complex, including Río Negro virus (RNV), have been shown to circulate widely in Argentina and neighboring countries. RNV has been detected in mosquitoes and rodents from multiple Argentine provinces, indicating sustained enzootic transmission and geographic expansion (4–8). Serological evidence of RNV infection has also been reported in humans and equines across several regions of Argentina and Uruguay, with equine seroprevalence reaching up to 20% in some areas (9,10). In addition, bats have been proposed as potential hosts in the natural cycle of RNV, following the detection of viral genomes in oral swabs (11,12). Importantly, the first clinical case of human infection with RNV was recently reported in Bolivia highlighting its capacity to cause human disease and underscoring the need for improved surveillance and diagnostic capacity. Altogether, these findings indicate that RNV represents a potential threat to both human and equine health in the Southern Cone of Latin America.

Epizootic strains of Venezuelan equine encephalitis virus (VEEV) have been extensively studied due to their capacity to cause severe outbreaks in equine and human populations. Experimental studies in laboratory mice have shown that these variants replicate systemically, invade the central nervous system, and trigger robust inflammatory responses that contribute to neurological disease (13–17). In contrast, the pathogenic potential of natural enzootic VEEV species not belonging to subtype I has been far less explored in animal models. Early studies using non-human primates or rodents infected with enzootic strains such as Mucambo virus or Everglades virus reported limited or mild disease following peripheral inoculation, despite evidence of viral neuroinvasion (18,19). With regard to Río Negro virus (RNV, subtype VI), early in vivo characterizations demonstrated an apparent lack of pathogenicity in immunocompetent adult mice and guinea pigs, even at high inoculation doses (20). However, experimental approaches restricted to adult immunocompetent models may overlook age- or immunity-dependent pathogenic features that emerge under alternative conditions (21–23). Consistent with this notion, young mice have been successfully used to reveal disease phenotypes caused by viruses that are otherwise non-pathogenic in adult wild-type animals, including alphaviruses such as Chikungunya virus (24).

The aim of the present study is to characterize the pathogenesis of RNV in a mouse model, with particular attention to the events underlying disease development. By analyzing the course of RNV infection in laboratory mice, we seek to expand the currently limited data available for evaluating the pathogenic and epidemic potential of this neglected alphavirus. In the context of ongoing ecological change, including landscape transformation and climate-driven shifts in vector ecology, understanding the pathogenic capacity of enzootic arboviruses such as RNV is of increasing relevance (25). Of particular concern is the potential involvement of enzootic or epizootic Venezuelan equine encephalitis virus strains in emergent urban transmission cycles, especially given the demonstrated competence of Aedes aegypti to transmit both epidemic and enzootic variants of the virus (26).

## Methods

### Virus and mice

The Rio Negro virus strain (RNV, strain F89) used in our study was originally isolated from rodents captured in Formosa province, Argentina, in 1991 (5), and underwent three passages in suckling mouse brains. This virus was provided by Dr. Marta S. Contigiani from the Laboratory of Arboviruses at the Virology Institute of the Universidad Nacional de Córdoba, Argentina. Working stocks were prepared directly from this material by homogenizing the previously infected mouse brain tissue (10% w/v) in Eagle’s minimum essential medium (MEM) supplemented with 10% fetal bovine serum and 1% gentamicin. Virus titers were determined using a Vero cell plaque assay and expressed as plaque forming units per milliliter (PFU/mL) (27). For all experiments, virus inoculations were performed via dorsal subcutaneous injection, administering 10³ PFU in 0.1 mL of MEM per mouse. Control groups were inoculated with 0.1 mL of MEM supplemented with 10% fetal bovine serum and 1% gentamicin. C57BL/6 mice were housed in a specific-pathogen-free (SPF) environment, kept in microisolated ventilated cages, and handled by highly trained personnel within a laminar BSL2 flow hood. All animal procedures were approved by the Institutional Committee for the Care and Use of Laboratory Animals (CICUAL) of the Facultad de Ciencias Médicas, Universidad Nacional de Córdoba, under protocol number V-14/2022. All *in vivo* and *in vitro* experiments with RNV were conducted under enhanced biosafety level 2 (BSL-2+) conditions, in compliance with institutional risk assessments and local regulations. Enhanced laboratory practices were applied throughout all procedures to minimize exposure risk.

### Survival

To determine the most appropriate age group for pathogenicity studies, we evaluated the susceptibility of C57BL/6 mice of different ages to RNV infection. Mice aged 16, 17, 18, 19 and 20 days were used. For each age group, animals from three different litters were included, resulting in groups of 20-24 mice, depending on litter size. After infection, survival was monitored daily for 15 days post-infection (dpi). The confirmation of infection in deceased animals was achieved by detecting infectious viral particles in brain homogenates using Vero cell plaque assays (27). Surviving mice were tested for neutralizing antibodies using the plaque reduction neutralization test (PRNT) (28) on sera samples (diluted at 1:5 in MEM).

### Viremia and viral load in tissues

Viremia was assessed in groups of 5 mice at different time points, expressed as hours post-infection (hpi): 12, 14, 19, 24, 30, 48, 72, and 96. Control groups consisted of three mock-infected (MEM) mice per time point. Blood samples were collected at each time point via submandibular vein puncture. After induction of anesthesia with isoflurane inhalation (2%), euthanasia was carried out via cervical dislocation by trained personnel. Viral loads in serum samples were determined using a standard Vero cell plaque assay and expressed as the logarithm of PFU per milliliter (log PFU/mL).

To evaluate viral dissemination, 27 mice aged 18 days were infected, and three animals were sacrificed daily from day 0 to 9 post-inoculation for tissue collection, including brain, spleen, axillary lymph nodes (near the inoculation site), lungs, pancreas, liver, heart, and thymus. In addition, 3 age-matched uninfected mice were used as controls and sacrificed on day 9 of the experiment. Prior to sacrifice, mice were anesthetized with an intraperitoneal (IP) injection of xylazine (10 mg/kg) and ketamine (100 mg/kg) and perfused transcardially with PBS until the liver blanched (approximately 4–5 minutes). Approximately half of each tissue sample was weighed, suspended in MEM supplemented with 10% FBS and 1% gentamicin (1/10 w/v), and homogenized using a sterile cold mortar and pestle. The homogenates were clarified by centrifugation at 10,000 g for 30 min, and the supernatants were used to determine viral loads.

### Histology

A fraction of each collected tissue sample was fixed overnight in 10% neutral buffered formalin, embedded in paraffin, sectioned (4 μm), and stained with hematoxylin and eosin (HE) for qualitative histologic examination. Micrographs were taken using a Leica ICC50 E camera attached to a Leica DM500 binocular microscope.

### Spleen immunofluorescence

Spleens from 2 control and 2 RNV-infected mice were collected at 2, 4 and 6 dpi and immediately flash-frozen in liquid nitrogen. Cryosections (7 µm) were fixed in cold acetone for 10 min, air-dried at room temperature (RT), and stored at −80 °C until further use. Prior to staining, slides were rehydrated in Tris buffer and incubated for 30 min at 25°C in blocking solution containing 10% normal mouse serum diluted in Tris buffer (29). After blocking, slides were incubated for 50 min at 25°C with dif ferent combinations of the following anti-mouse Abs: PE-labeled anti-CD8a (Cat# 12-0081-83, clone 53-6.7, dilution 1/100), APC-labeled anti-B220 (Cat# 17-0452-83, clone RA3-6B2, dilution 1/200) from eBiosciences, and AlexaFluor 488-labeled anti-CD169 (Cat# 142419, clone 3D6.112, dilution 1/150) from Biolegend. Slides were mounted using FluorSave reagent (Merck Millipore), and images were acquired with an Olympus FV1000 confocal microscope. To ensure the representativeness of the qualitative findings, at least five sections from different regions of each spleen were examined, and multiple follicles per section were analyzed to confirm the consistency of the observed histological patterns. Image analysis was performed using ImageJ64 v1.52e (NIH, USA).

### Analysis of neurodegeneration

Three infected mice were sacrificed at 2 dpi (n=3, with no signs of illness) or at 8 dpi (n=3, with neurological signs). Chloral hydrate (0.3g/Kg) was injected intraperitoneally (IP) for anesthesia and each mouse was transcardially perfused, rinsed with glucose (0.4%), sucrose (0.8%), and sodium chloride (0.8%), and fixed with 4% paraformaldehyde (PFA) in 0.2 M borate buffer (pH 7.4). After immersion in 30% sucrose, brains were cut in frontal and sagittal sections (40µm) using a cryostat. A subset of sections was reserved for immunohistochemical analyses. Neurodegeneration was assessed in brain tissue using the amino-cupric-silver (A-Cu-Ag) staining technique, performed according to the method described by de Olmos et al. (30). Briefly, brain sections were rinsed with Milli-Q water and incubated in a silver nitrate solution at 50 °C (pre-impregnation). Once back at room temperature, sections were rinsed with acetone and transferred to a concentrate silver diamine solution for 40 min. They were immersed in a formaldehyde/citric acid solution for 25 min and the reaction was stopped in 0.5% acetic acid. A two steps bleaching was performed to remove any non-specific deposit of silver, first in 6% potassium ferricyanide, washed with Milli-Q water, then transferred to 0.06% potassium permanganate for 20s. After washing, stabilization was achieved in 2% sodium thiosulphate, then the sections were washed again, transferred to a fixer solution for 1 min, and mounted on slides. Once dried, slides were cleared by immersion in xylene for 10 min before coverslipping. Images were obtained in an optic microscope (OlympusCX40) equipped with a Zeiss AxioCam ERc5s camera. Neuronal degeneration was evaluated qualitatively on cortical structures and striatum of infected mice. Sections of uninfected mice were analyzed as controls.

### Brain immunohistochemistry

Brain sections (40 µm), obtained from the same perfused and cryoprotected brains described above for neurodegeneration analysis, were processed for free-floating immunofluorescence in 6-well plates under constant agitation, following standard protocols. Briefly, 10–15 sections per mouse were washed three times for 5 min each in PBS. Sections were then blocked and permeabilized in PBS containing 0.3% Triton X-100 and 3% FBS for 2 hours at RT. Primary antibody incubation was performed for 24 h at 4 °C under agitation, using rabbit polyclonal anti-glial fibrillary acidic protein (GFAP; 1:500; cat. no. G3893, Sigma-Aldrich) diluted in blocking solution (PBS with 1% FBS and 0.3% Triton X-100). After washing, sections were incubated for 1 hour at RT with Alexa Fluor 488-conjugated donkey anti-rabbit IgG (1:1000; cat. no. A21206, Invitrogen), followed by counterstaining with DAPI in PBS for 15 min. Finally, sections were washed three times in PBS, mounted on gelatin-coated slides with Mowiol, coverslipped, and dried overnight. Images were acquired using a confocal fluorescence microscope (Olympus FV1200, Japan).

### Detection of inflammatory cytokines

Two groups of three infected mice were euthanized at 2 dpi (first group) or immediately after the onset of neurological symptoms (6 dpi). An uninfected control group (n=3) was processed in parallel. Brains were collected, flash-frozen in liquid nitrogen, and stored at −80°C. After weighing and thawing, tissues were minced using two scalpels and processed according to the protein isolation method described by Amsen et al. (31), keeping all samples on ice throughout the procedure. Briefly, tissues were homogenized using a tissue mixer (Janke & Kunkel) in a lysis buffer lacking SDS. Lysates were then sonicated using a probe sonicator (five 30-s pulses with 30-s intervals) and centrifuged at 14,000 × g for 20 min at 4°C. Supernatants were collected and cytokine levels were measured using a multiplex Cytometric Bead Array (CBA) system, following the manufacturer’s instructions (Mouse Inflammation Kit, BD Biosciences). Standard and test samples (infected and control groups) were analyzed using an Accuri C6 flow cytometer (BD).

### Statistical analyses

Data were analyzed using GraphPad Prism (version 8.0.2). Normality was assessed with the Shapiro-Wilk test, followed by a one-way ANOVA to compare mean values across experimental groups or time points. Post-hoc comparisons were performed using Tukey’s multiple comparisons test, with statistical significance set at p < 0.05. Survival data were analyzed using Kaplan–Meier survival curves. Deaths were coded as events (value = 1), and animals surviving until the end of the observation period were treated as censored data (value = 0). Differences among age groups were assessed using the log-rank (Mantel–Cox) test, and age-dependent trends in survival were evaluated using the log-rank test for trend. Median survival times (in dpi) were calculated for each age group. For viremia, viral titers were directly analyzed without prior adjustment and are presented as the logarithm of plaque-forming units per milliliter (log PFU/mL). For organotropism, viral titers in tissue homogenates collected at different dpi were analyzed using the same statistical approach to evaluate differences among time points. For cytokine quantification by CBA, raw cytometry data were first processed using a nonlinear regression 3PL curve fit to obtain interpolated values, which were then subjected to statistical analysis as described.

## Results

### RNV infection causes age-dependent disease and mortality in mice

To determine an appropriate murine model for studying RNV pathogenesis, we first assessed age-dependent susceptibility in infant C57BL/6 mice. Animals aged 16 to 20 days were subcutaneously infected and monitored for signs of illness, mortality, and evidence of infection. Marked differences in survival were observed among age groups (**Fig 1)**. The log-rank (Mantel-Cox) test revealed highly significant differences between survival curves (χ² = 86.49, df = 4, p < 0.0001), indicating that survival probability strongly depended on age at the time of infection. The log-rank test for trend also showed a significant linear association (χ² = 57.83, df = 1, p < 0.0001), consistent with a progressive increase in survival with age. All 20-day-old mice survived without apparent illness, unlike younger groups that showed 100% mortality at 16–18 days of age and early clinical signs (ruffled fur, lethargy) appearing 2–3 dpi (**Fig 1**). In these younger animals, the disease evolved into ataxia and paralysis before culminating in death. Infection was confirmed in all deceased mice by detecting infectious viral particles in brain homogenates, whereas surviving animals developed neutralizing antibodies against RNV, with titers exceeding 1:20.

**Fig 1.**
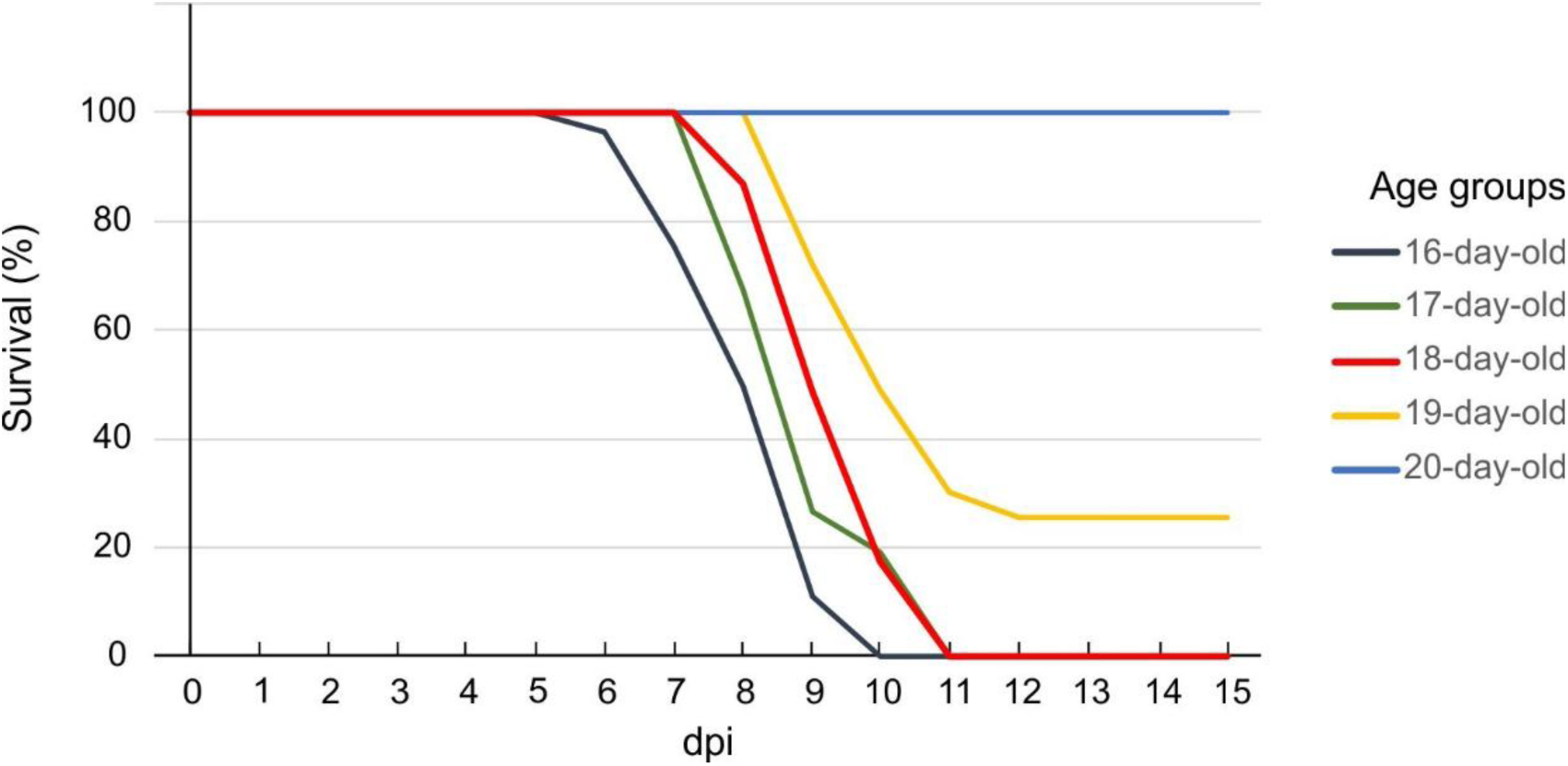
Age-dependent survival following RNV infection in C57BL/6 mice. Infant mice aged 16-20 days were infected and monitored daily for survival over a 15-day period (dpi, days post-infection). Each curve represents pooled data from three litters per age group (n = 20–24 per group).

The survival curves clearly illustrate a sharp decline in susceptibility to RNV infection between 18 and 20 days of age. The 18-day-old group, being the oldest cohort to exhibit complete mortality, was therefore selected for subsequent pathogenicity experiments.

### RNV infection in infant mice results in systemic dissemination and broad organ tropism

Viremia was detected between 14- and 48-hours hpi, with a peak of 2.85 ± 0.03 log₁₀ PFU/mL at 30 hpi. No secondary viremia was detected in blood samples collected daily up to 10 dpi, and no infectious virus was detected in serum from mock-infected control mice at any time point (**Fig 2**).

**Fig 2.**
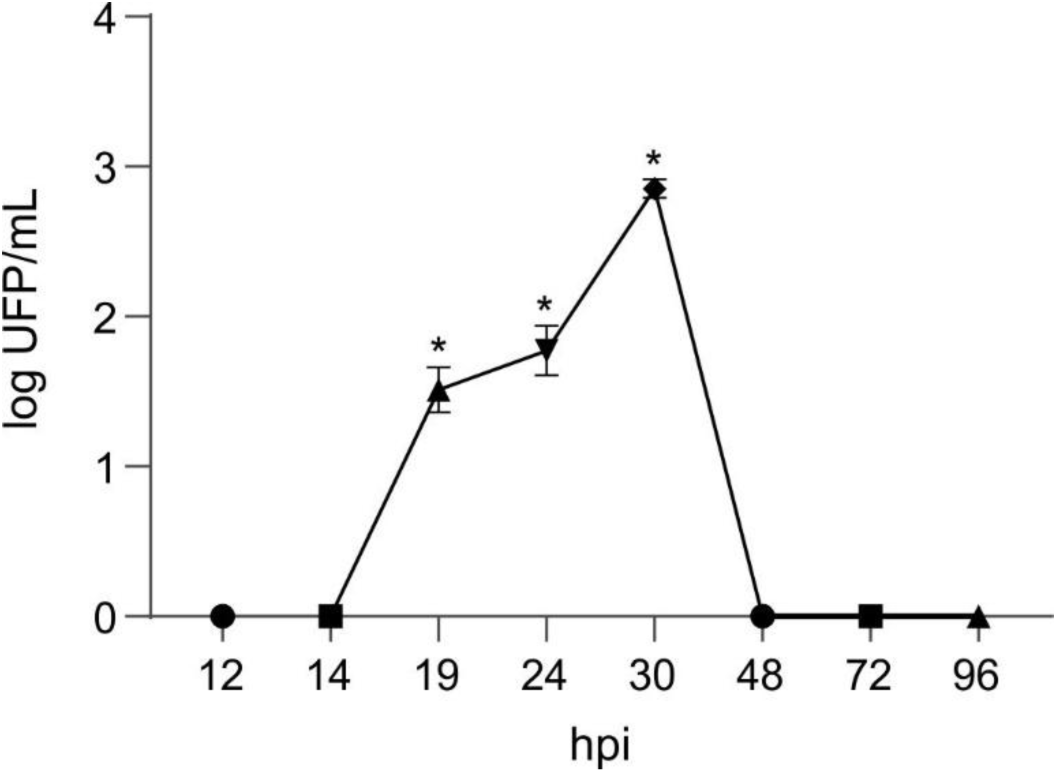
RNV infection results in transient viremia in infant mice. Viral titers in serum samples are expressed as log PFU/mL at different hours post-infection (hpi). Data points represent the mean of five mice per time point, with error bars indicating standard deviations. Asterisks (*) denote statistically significant increases in mean viral titers between consecutive time points (p < 0.05, one-way ANOVA followed by Tukey’s multiple comparisons test).

To further investigate the systemic impact of infection, we analyzed the distribution of viral particles across multiple organs. No infectious virus was detected in the heart or liver, whereas significant viral replication was observed in the spleen, lymph nodes, thymus, pancreas, lung, and brain (**Fig 3**). Viral dissemination occurred rapidly, with detectable loads in lymphatic tissues as early as 24 hpi. Peak mean viral titers occurred on day 2 post-infection (pi) in the spleen (5.11 ± 0.45 log PFU/g) and lymph nodes (4.38 ± 0.38), on day 4 pi in the pancreas (4.88 ± 0.17), lungs (3.16 ± 0.72), and thymus (5.18 ± 0.33), and on day 5 pi in the brain (7.70 ± 0.10). A one-way ANOVA revealed significant differences in viral titers across tissues (F = 40, p < 0.05). Post hoc comparisons showed that viral loads in the brain were significantly higher than in all other tissues (p < 0.05). Moreover, brain infection persisted the longest, extending up to day 9 pi, coinciding with the terminal phase of neurological disease.

**Fig 3.**
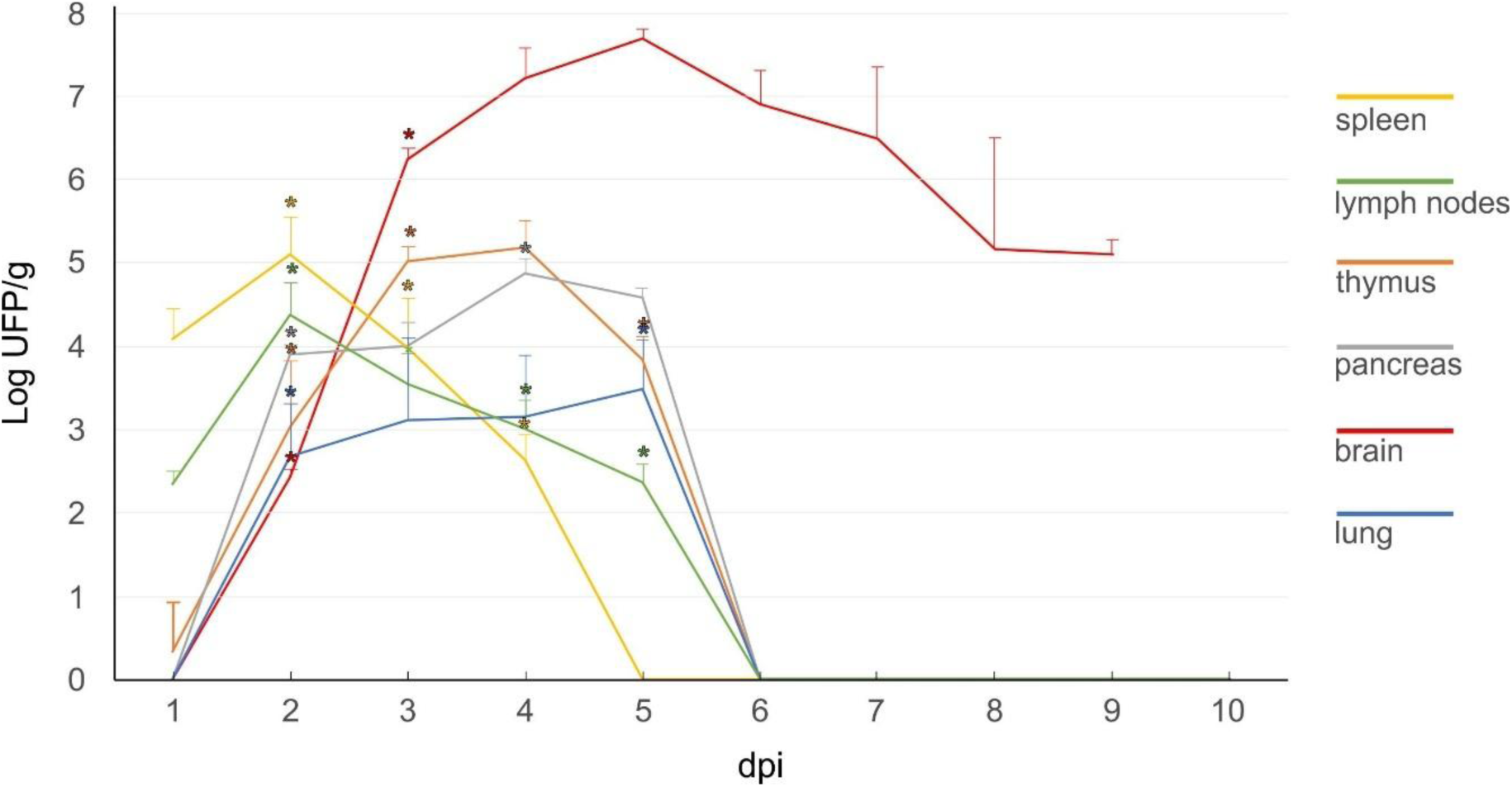
RNV infection leads to systemic dissemination and broad tissue tropism. Viral titers (log PFU/g) were measured in various organs at different days post-infection (dpi) using Vero cell plaque assays. Data points represent the mean viral load per gram of tissue from three mice per time point, with error bars indicating standard deviations. Asterisks (*) denote statistically significant differences in viral titers between consecutive time points within each organ (p < 0.05, one-way ANOVA followed by Tukey’s multiple comparisons test).

### RNV induces histopathological alterations in infected tissues

We qualitatively assessed the effects of RNV infection on selected tissues at different dpi, corresponding to periods of active viral replication. In HE-stained spleen sections from infected mice, changes suggestive of architectural disorganization were observed by 6 dpi, with the boundaries between white and red pulp appearing less clearly defined (**Fig 4C-D**).

**Fig 4.**
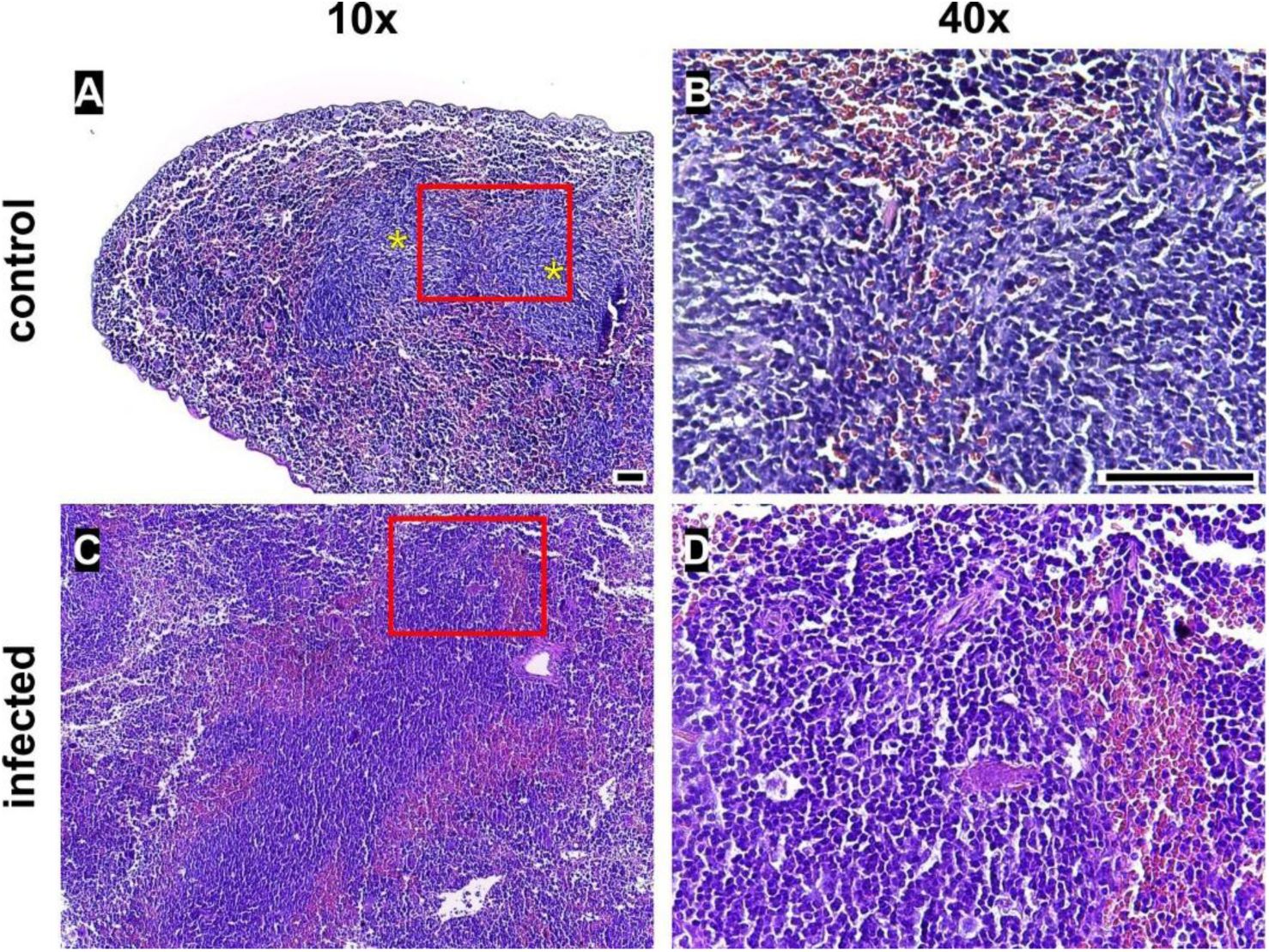
RNV infection induces alterations in splenic architecture. Representative HE-stained spleen sections (4 µm) at 10x (left panels) and 40x (right panels) magnification. (A-B) Mock-infected control mouse showing preserved splenic architecture, with well defined white pulp follicles (yellow asterisks). (C-D) Spleen from a RNV-infected mouse at 6 day post-infection. Magnification: 10x (A, C) and 40x (B, D). Scale bars: 500 µm (10x and 40x). Red boxes in 10x panels indicate the region shown at higher magnification on the right. Images are representative of one out of 3 mice per group.

To further investigate the splenic alterations revealed by HE staining, we performed immunofluorescence to evaluate the organization of immune cell compartments. In uninfected control mice, the spleen showed a well-organized structure with clearly defined B-cell (B220^+^) and T-cell (CD8^+^) zones, and a discrete presence of CD169^+^ metallophilic macrophages (**Fig 5A**). In contrast, infected mice exhibited a severe and progressive disorganization of the lymphoid microenvironment. This pattern was consistently observed across all analyzed sections, where as early as 2 dpi, B-cell follicle boundaries became diffuse, with notable infiltration of CD8^+^ T cells into B220^+^ areas (**Fig 5B**). By 6 dpi, this disorganization was markedly exacerbated, with widespread loss of compartmentalization and increased dispersion of both CD8^+^ T cells and CD169^+^ metallophilic macrophages across the tissue (**Fig 5C**).

**Fig 5.**
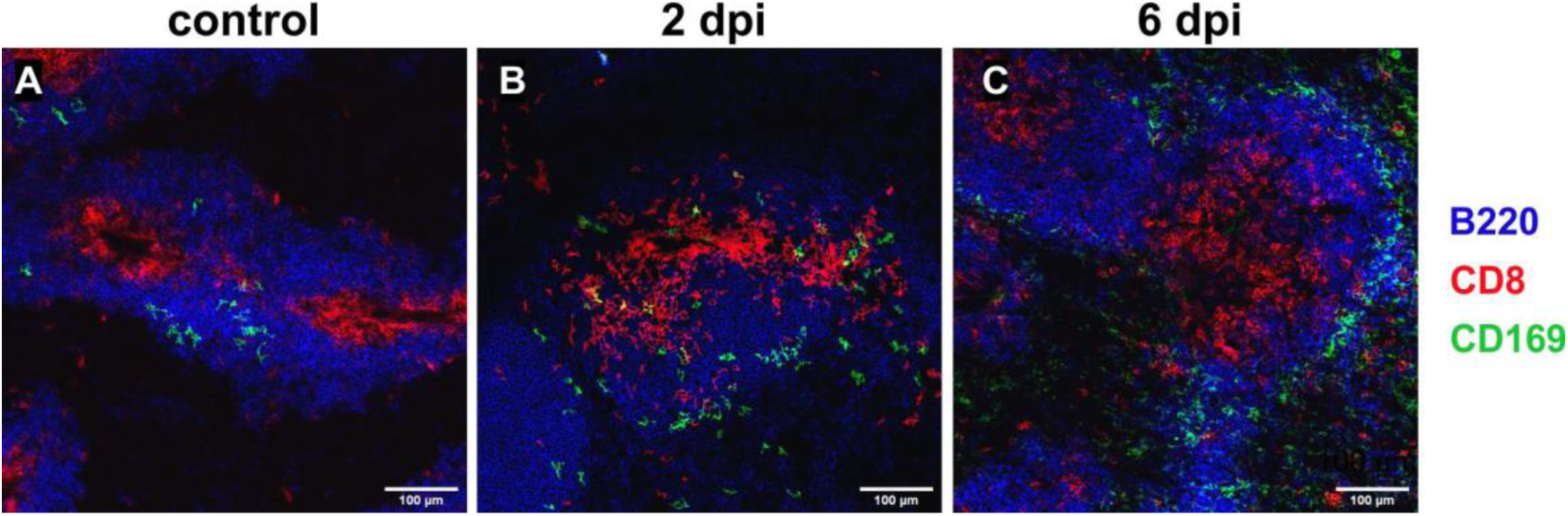
Disruption of splenic immune architecture in RNV-infected mice. Representative images of spleen sections (7 µm) from uninfected control mice **(A)** and RNV-infected mice at 2 **(B)** and 6 **(C)** days post-infection (dpi), stained with anti B220 (blue, B cells), anti CD8 (red, T cells), and anti CD169 (green, macrophages). Images were acquired with an Olympus FV1000 confocal microscope at 20x magnification. Scale bars: 100 µm. Findings are representative of multiple non-consecutive sections analyzed per animal (n = 2 mice per group/time point).

HE-stained lung sections from uninfected mice showed preserved alveolar architecture, with thin septa and sparse interstitial cellularity (**Fig 6A–B**). In contrast, infected mice analyzed at 8 dpi displayed pronounced thickening of alveolar septa and a marked increase in interstitial cellularity, consistent with inflammation and tissue remodeling (**Fig 6C and E**, rectangles; **Fig 6D and F**, asterisks).

**Fig 6.**
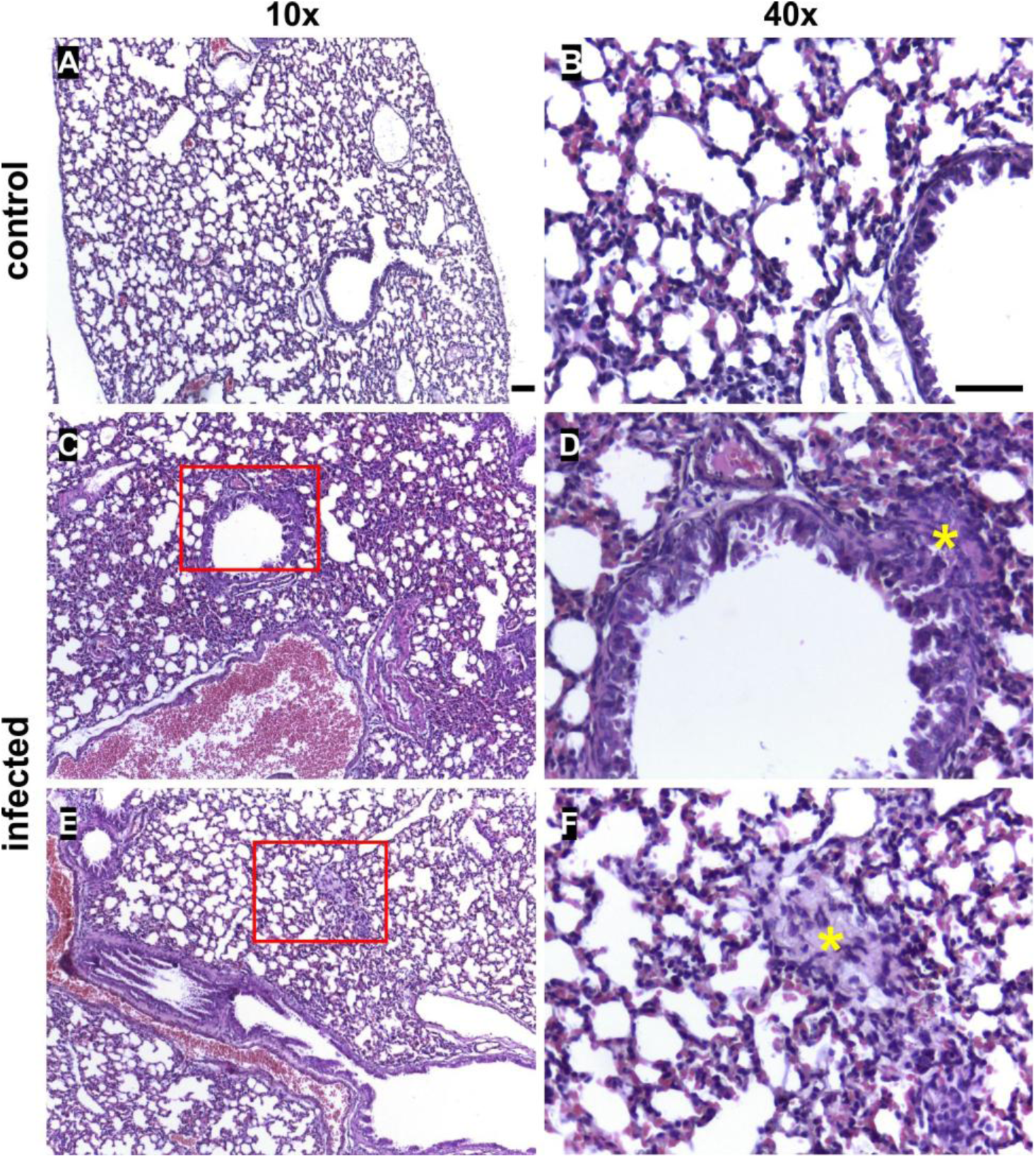
RNV infection induces inflammatory pathology in the lungs. HE-stained lung sections (4 µm) from uninfected control mice (A-B) and RNV-infected mice at 8 days post-infection (C-F). Panels A, C, and E were captured at 10x magnification; panels B, D, and F at 40x magnification. Magnification: 10x (A, C, E) and 40x (B, D, F). Scale bars: 500 µm (10x and 40x). Red rectangles in the 10x images indicate the areas shown at higher magnification in the corresponding 40x magnification. Photographs are representative of one out of 3 mice at each time point.

Consistent with the neurological symptoms observed in infected animals, clear pathological alterations were also detected in brain sections from RNV-infected mice, which became more pronounced at later stages of infection. HE-stained sections revealed localized inflammatory processes in the meninges, characterized by edema and immune cell infiltration (**Fig 7C**, rectangle; **Fig 7D**, arrow). In addition, multiple foci of cell infiltration were found throughout the cortex, frequently coalescing into discrete clusters (**Fig 7E**, rectangle; **Fig 7F**, asterisks). In addition, HE sections of the olfactory bulb from infected mice revealed vascular congestion with intraluminal nucleated cells, indicative of local inflammation (**Fig 8C and D**, arrows).

**Fig 7.**
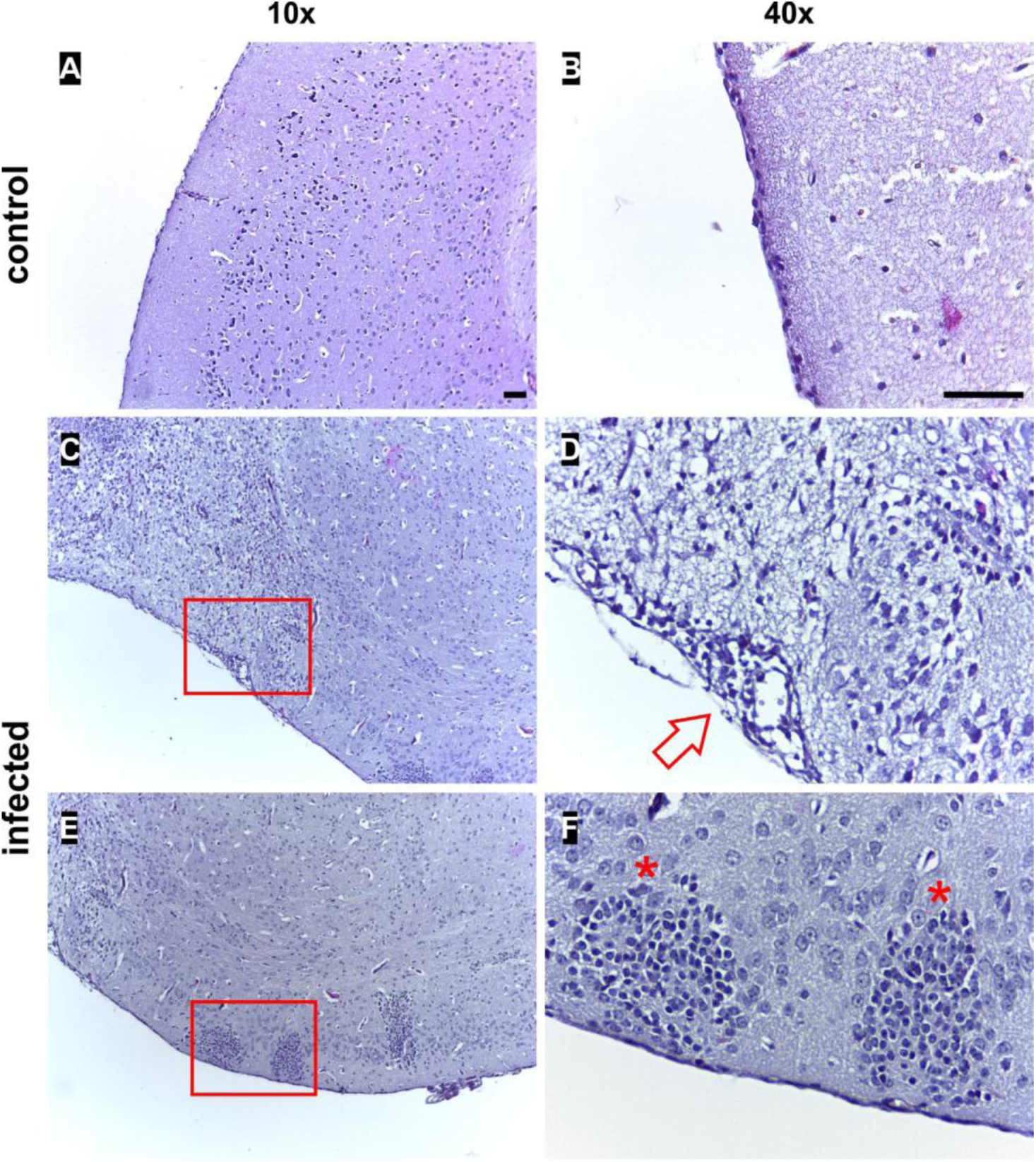
RNV infection results in histopathological changes in the brain. Panels A-B correspond to uninfected control mice. Panels C-F show brains from infected mice at 8 days post-infection. Sections were stained with HE and cut at 4 µm thickness. Magnification: 10x (A, C, E) and 40x (B, D, F). Red rectangles in the 10x images indicate the areas shown at higher magnification in the corresponding 40x magnification Scale bars: 500 µm (10x and 40x). Photographs are representative of one out of 3 mice at each time point.

**Fig 8.**
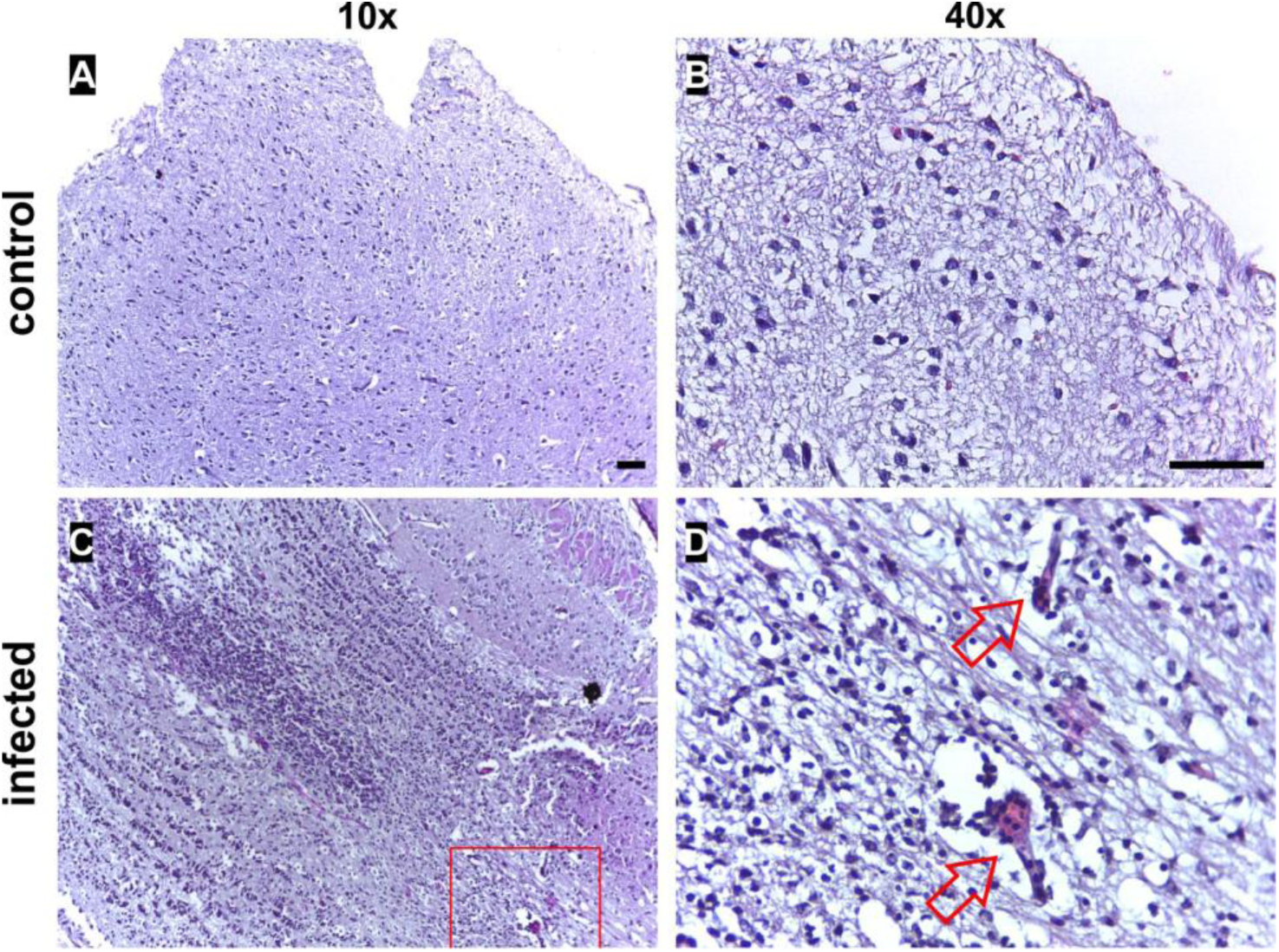
Histopathological alterations in the olfactory bulb following RNV infection. Panels A-B correspond to uninfected controls. Panels C-D show infected mice at 8 days post-infection. Sections were stained with HE and cut at 4 µm thickness. Magnification: 10x (A, C, E) and 40x (B, D, F). The red rectangle in the 10x image indicate the area shown at higher magnification in the corresponding 40x magnification. Scale bars: 500 µm (10x and 40x). Images are representative of three mice per group.

### RNV infection triggers neurodegeneration and glial activation

We subsequently investigated whether RNV infection could induce neurodegeneration in specific brain regions, both prior to the onset of neurological symptoms (2 dpi) and at terminal stages of disease (8 dpi). **Fig 9** shows sagittal brain sections stained with the A-Cu-Ag technique, highlighting the progression of degeneration. As a reference, uninfected mice displayed only isolated foci of physiological cell death, likely associated with normal postnatal development (**Fig 9A-B**). At 2 dpi, infected mice already harbored high viral loads in the brain (exceeding 6 Log PFU/g), but only sparse silver-stained foci were observed (**Fig 9C-D**). In contrast, the brains of symptomatic animals at day 8 pi exhibited widespread somatodendritic and terminal degeneration, prominently clustered around blood vessels (**Fig 9E-F**).

**Fig 9.**
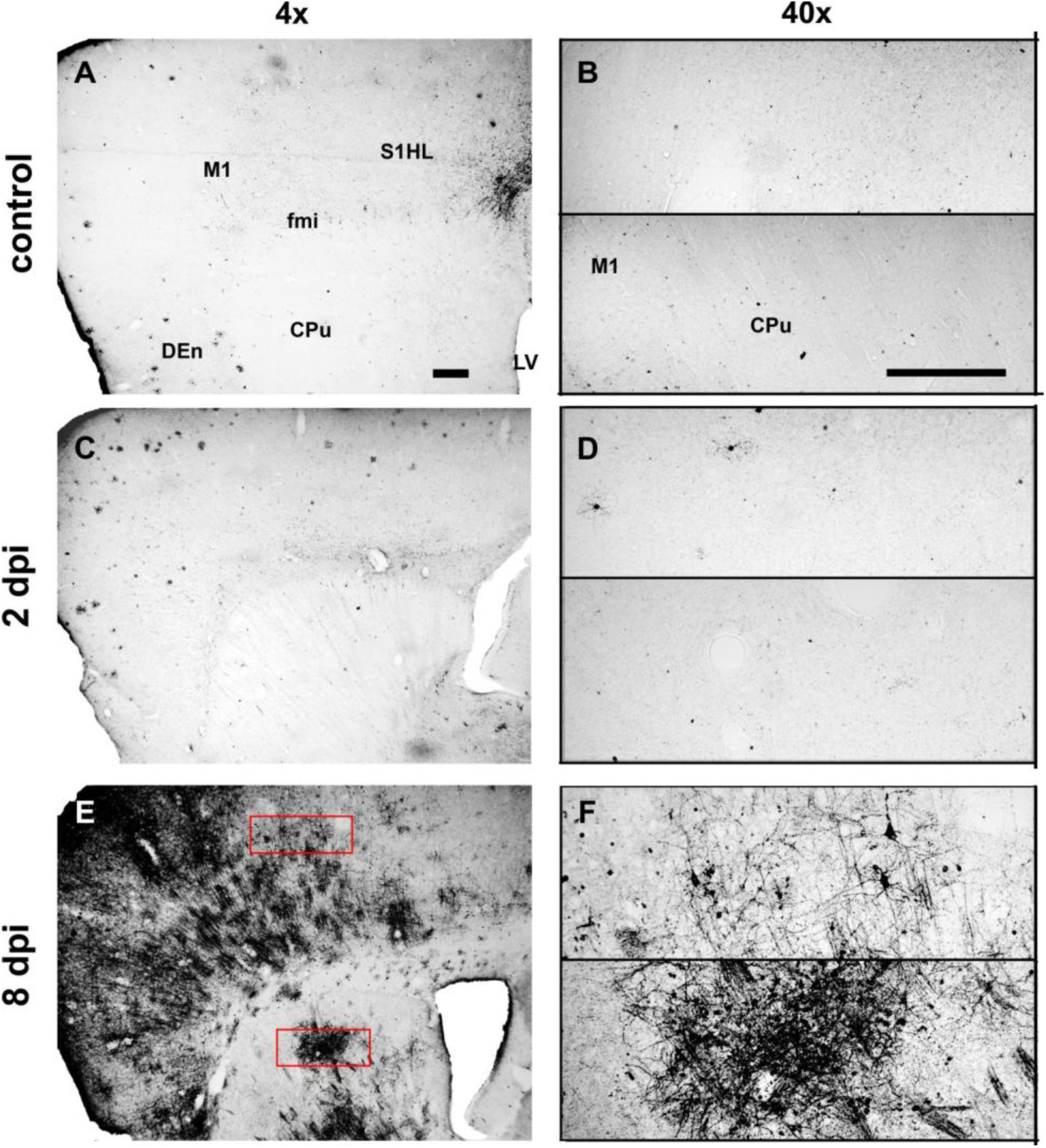
RNV infection induces neurodegeneration in the brain. Representative sagittal brain sections stained with the amino-cupric-silver (A-Cu-Ag) technique. Panels A and B show an uninfected age-matched control mouse. Panels C and D correspond to mice infected with RNV and euthanized at 2 days post-infection (dpi). Panels E and F display brain sections from infected mice at 8 dpi. Degenerative foci appear as dark silver-stained structures. Left panels: 4x magnification; right panels: 40x magnification. M1: primary motor cortex; S1HL: somatosensory cortex (hind limb area); CPu: caudate-putamen; fmi: forceps minor of corpus callosum; DEn: dorsal endopiriform nucleus; LV: lateral ventricle. Scale bars: 500 µm (4x) and 100 µm (40x). Images are representative of 3 mice per group.

While infected mice already harbored high viral loads in the brain at 2 dpi (exceeding 6 Log PFU/g), only sparse silver-stained foci were observed at this stage (**Fig 9C-D**). In contrast, the brains of symptomatic animals at day 8 pi exhibited widespread somatodendritic and terminal degeneration, prominently clustered around blood vessels (**Fig 9E-F**). As a reference, uninfected mice displayed only isolated foci of physiological cell death, likely associated with normal postnatal development (**Fig 9A-B**).

To further investigate the neuroinflammatory response to RNV infection, we performed a qualitative analysis of astrocyte activation using anti-GFAP staining on brain sections. The staining was performed on coronal sections encompassing the entire brain, and GFAP signal alterations were observed in multiple regions, including the hippocampus, cerebral cortex, and cerebellum. However, for consistency and image clarity, representative visualization was focused on the hippocampal region, where staining was most robust and reproducible across samples. As shown in **Fig 10**, control mice (**Fig 10A-C**) exhibited sparse and finely branched GFAP^+^ astrocytes, mostly confined to specific hippocampal layers. In contrast, infected mice (**Fig 10D-F**) showed progressive increases in GFAP signal intensity and astrocyte hypertrophy. At 2 dpi, GFAP^+^ processes appeared more numerous and thicker, suggesting early astrocyte activation. These changes were markedly intensified at 8 dpi, with dense GFAP immunoreactivity and enlarged astrocytic processes, consistent with a robust astrogliosis. While the temporal and spatial overlap between gliosis and neuronal injury remains to be fully resolved, the presence of marked astrocyte activation in multiple brain areas supports a widespread neuroinflammatory response to RNV infection.

**Fig 10.**
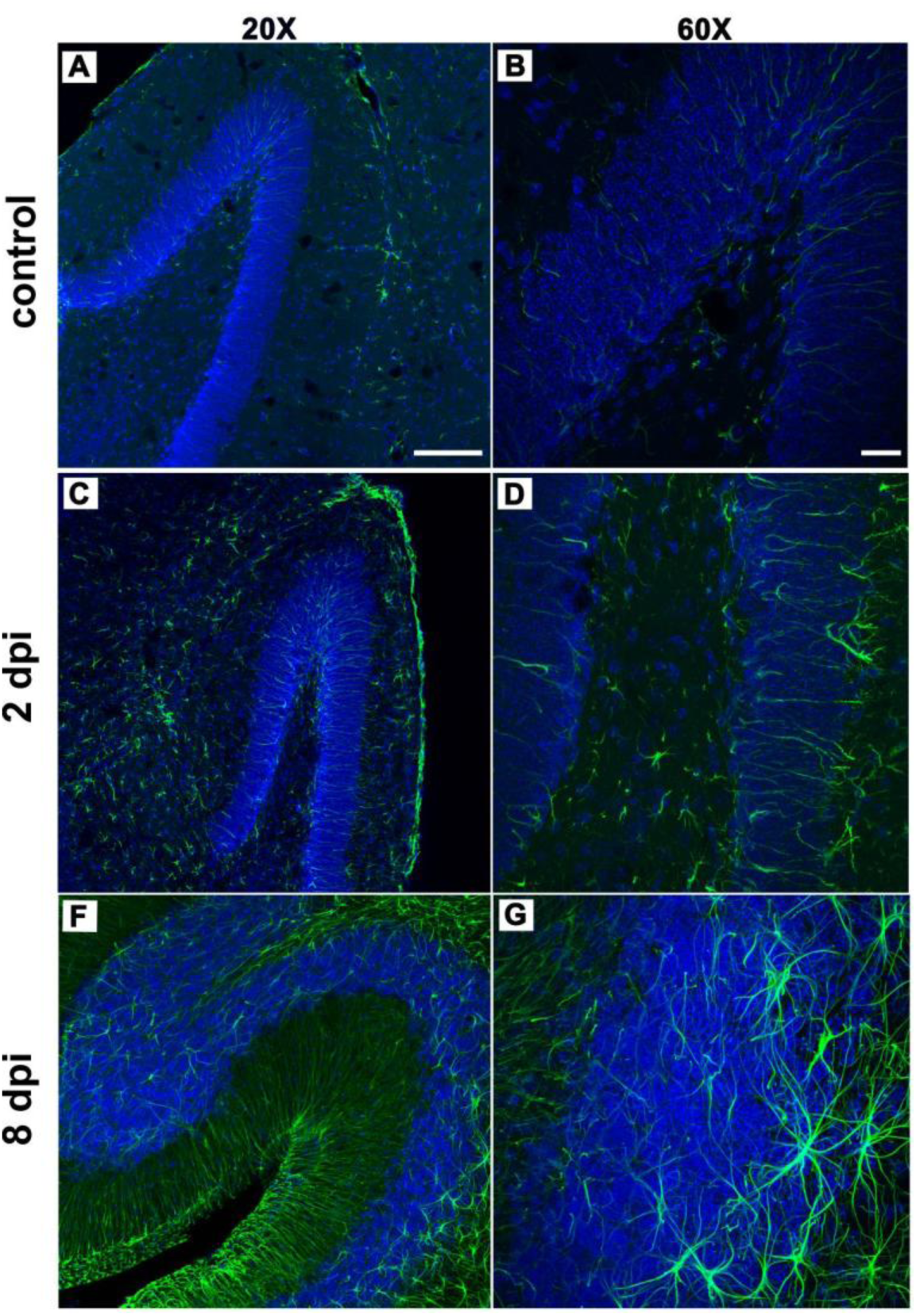
Astrocyte activation in the hippocampus during RNV infection. Immunofluorescent staining of sagittal brain sections at the hippocampal level using anti-GFAP (green) and DAPI (blue). Panels A-B correspond to uninfected controls. Panels C-D show infected mice at 2 days post-infection (dpi). Panels E-F show infected mice at 8 dpi. Left panels: 20x magnification; right panels: 60x magnification. Scale bars: 100 µm (20x) and 20 µm(60x). Images are representative of 3 mice per group.

### RNV induces pro-inflammatory cytokines in the brain

We next sought to determine whether this glial response was associated with increased expression of inflammatory mediators. To this end, we quantified inflammatory cytokines in the brain using a commercial CBA. A panel of six cytokines was analyzed (IL-6, IL-10, MCP-1, IFN-𝛾, TNF, IL-12p70) in brains collected at 2 dpi and at 6 dpi. As shown in **Fig 11**, symptomatic animals exhibited significantly elevated levels of IFN-𝛾, IL-6, and MCP-1 compared to both control and 2 dpi groups. Although no statistically significant differences were detected for TNF or IL-10, an increased dispersion of values was observed in symptomatic animals. IL-12p70 levels remained undetectable across experimental groups. These results indicate that the onset of neurological symptoms is associated with a robust local induction of key pro-inflammatory mediators in the brain.

**Fig 11.**
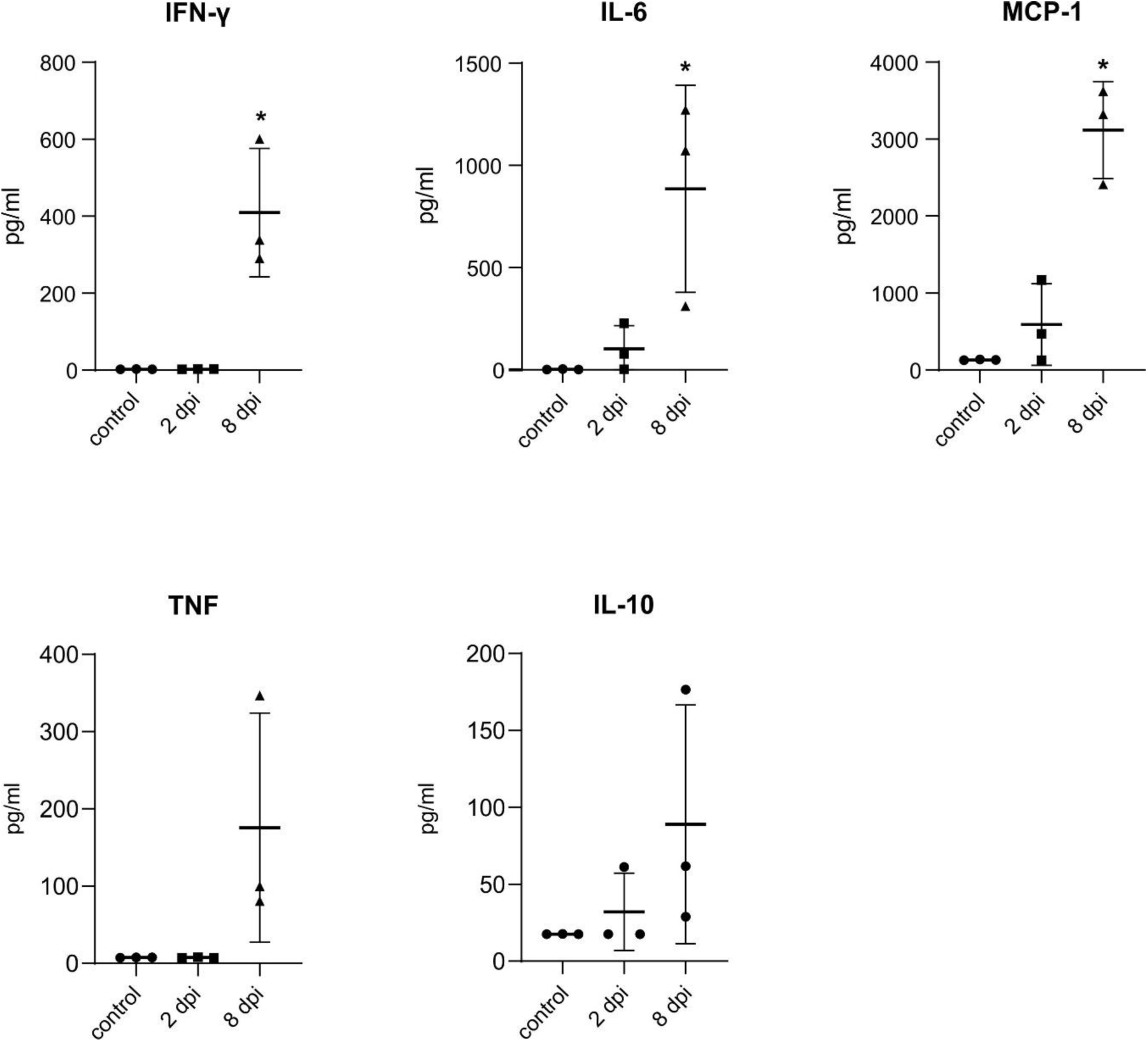
RNV infection induces inflammatory cytokine responses in brain tissue. Levels of IFN-𝛾, IL-6, MCP-1, TNF, and IL-10 were quantified using a multiplex Cytometric Bead Array (CBA) system. Experimental groups included uninfected controls (n=3), mice at 2 days post-infection (dpi) (n=3), and at 6 dpi (n=3). Data are presented as individual values with group means ± SEM. Statistical comparisons were performed using one-way ANOVA followed by Tukey’s multiple comparisons test for IFN-𝛾, IL-6, and MCP-1, with significant differences indicated by * (p < 0.05). For IL-10 and TNF, no significant differences were found (p = 0.2255 and p = 0.0841, respectively). These results are from a single experiment using three animals per group.

## Discussion

In general, animals infected with enzootic VEEV subtypes do not develop detectable viremia or disease (32), and virulent phenotypes in epizootic strains have been associated with specific sequences variants in the viral envelope glycoproteins (33). Although RNV has not been associated with large outbreaks of acute illness, its silent circulation may pose a potential - or underrecognized - risk to human and animal health, particularly in regions where enzootic transmission networks persist. The recently reported case of RNV-associated illness in an Argentine agricultural worker in Bolivia (34) highlights the potential of this virus to cause disease in humans, especially among those whose daily activities bring them into contact with disrupted transmission ecologies shaped by anthropogenic pressures.

In this context, we describe RNV pathogenesis in a susceptible mouse model. While previous studies reported no disease in adult immunocompetent laboratory rodents, we observed clear signs of progressive neurological illness and 100% lethality in infant 18-day-old wild-type mice. This pre-weaning stage represents a window of biological vulnerability that reflects the clinical observations in human VEEV-complex outbreaks, where children and older adults are disproportionately at risk for severe neurological disease, including convulsions, coma, and death (35,36). Importantly, the use of wild-type mice at this stage allowed us to investigate disease progression under conditions of natural immune immaturity, without the need for immunosuppressed or genetically modified models. Age-dependent susceptibility has been described in several viral models, including both arthritogenic and encephalitic alphaviruses, which tend to show increased tissue invasion and severity in younger hosts. (37–42).

To further characterize the pathogenesis of RNV in this model, we analyzed the profile and implications of viremia. Infection of infant mice with RNV resulted in a short-duration viremia with a relatively low peak viral titer (10^3^ PFU/mL) (**Fig 2**). Although no data are currently available for viremias induced by other strains within VEEV subtype VI, reports on related variants offer useful comparisons. Ludwig et al. (43) documented brief and low-level viremias (around 200 PFU/mL) in mice inoculated with attenuated VEEV strains developed for vaccine use, contrasting with epidemic strains such as Trinidad Donkey, which reached titers around 10^6^ PFU/mL. Notably, relatively low viremias not necessarily preclude transmission: mosquitoes - particularly of the genus *Culex* - have been shown to acquire enzootic VEEV strains from vertebrate hosts with viremias as low as 1,000-5,000 PFU/mL (44,45). Similar viremia levels were observed by Carrara et al. in *Proechimys chrysaeolus* spiny rats (mean peak: 3.3 log10 PFU/mL) experimentally infected with a subtype ID strain of VEEV (46). These rodents, considered natural reservoir hosts in endemic areas, showed no overt disease despite systemic infection. By contrast, our model revealed that even a modest and transient viremia was sufficient to support viral dissemination into multiple tissues, including the central nervous system. The absence of secondary viremia following the initial peak suggests that viral replication within target organs progressed independently of sustained viral presence in the bloodstream. Additionally, studies on enzootic subtype ID strains have demonstrated that low-level viremias in natural reservoir hosts can still support vector transmission and maintain enzootic cycles (47). Notably, despite comparable viremia levels, RNV infection led to efficient neuroinvasion and fatal encephalitis in infant mice.

To understand how RNV progresses following its brief viremic phase, we examined its distribution across multiple tissues. In our study, RNV reached high viral loads in spleens and lymph nodes as early as day 1 post-infection. Similar early replication in lymphoid tissues has been reported in adult mice infected with wild-type epizootic VEEV, where these compartments -draining lymph nodes and spleen- likely serve as primary sites of viral replication as early as 6 hpi (16). The spleen, as a major lymphoid filter and site of intense viral replication, may foster early quasispecies diversification, ensuring a variant spectrum broad enough to support viral tropism toward multiple tissues (48). While viral loads in peripheral organs declined after peaking, infection in the brain persisted throughout the course of neurological disease. The sustained presence of RNV in the brain, despite clearance from the bloodstream, suggests that CNS invasion occurs early during infection. Notably, the absence of detectable virus in other peripheral sites, such as the heart and liver, contrasts with the high titers observed in the infected tissues. While we cannot rule out the presence of viral RNA or low-level transient replication in these organs, our results indicate that they do not support robust productive infection under the conditions of this model. This organ-specific pattern of infection, particularly the sustained neurotropism, prompted us to further examine the histopathological alterations underlying disease progression.

Here, we report that RNV induced splenic architectural disorganization. The altered spatial distribution of B220+ B cells, CD8+ T cells, and CD169+ macrophages, was strikingly consistent across all examined specimens, showing a reproducible pattern of inflammation-associated remodeling (**Fig 5**). Proper spatial segregation of immune cell subsets is essential for adaptive immune responses, including antigen presentation and germinal center formation (49); thus, the extensive disruption observed here could be linked to impaired humoral immunity and viral persistence (50,51). In our model, the severity and consistency of this splenic collapse may help explain the failure of infant mice to mount an effective immune response, ultimately permitting uncontrolled viral dissemination, as evidenced by the development of pronounced pulmonary pathology.

Although moderate to severe interstitial pneumonia -characterized by infiltration of alveolar septa by neutrophils, lymphocytes, and macrophages- has been described in human VEEV infections (53), pulmonary pathology is not typically emphasized in experimental models. In contrast, in our model, RNV-infected mice exhibited overt signs of lung inflammation, including cellular infiltrates consistent with developing pneumonia (**Fig 6**), revealing the value of our model for investigating viral target cells in the lung. Notably, in the only confirmed human case of RNV infection to date, bibasilar crackles were reported during clinical examination (34), which may indicate lower respiratory tract inflammation. These findings raise the possibility that pulmonary involvement may be a more relevant feature of RNV pathogenesis than previously appreciated, particularly in naturally exposed or vulnerable hosts.

Following early dissemination and the establishment of viral replication in peripheral tissues, the progression of RNV infection in this model appears to exploit successive immunological vulnerabilities, enabling its spread toward organs that may provide permissive environments for replication. This organ-selective trajectory ultimately converges in the CNS, where viral replication coincides with a neuroinflammatory response whose intensity and spatial dynamics suggest a transition from protective immunity to immunopathology. Infectious virus in brain homogenates peaked at 5 dpi (**Fig 3**), temporally situating astrocytic activation within a phase of active viral replication in the CNS. This timing is consistent with reports for VEEV, in which GFAP upregulation emerges within 24-48 hpi, peaks at 3-5 dpi, and coincides with maximal viral replication and the onset of neurological symptoms (54). In epizootic VEEV strains, astrocytes have been identified as direct targets of infection through co-localization of viral antigens with GFAP *in vivo*. Furthermore, *in vitro* studies using primary or cultured astrocytes show virus-induced morphological alterations characteristic of reactive gliosis (55). Although we did not perform co-localization analyses or quantitative morphometric assessments in this study, the concurrent detection of elevated viral titers and pronounced astrogliosis, a common denominator of neurotropic viral infections, suggests a potential astrocytic contribution to the observed neuroinflammatory environment. Astrocytes are confirmed to be susceptible to infection by members of the Togaviridae family and are key mediators of the innate immune response via cytokine and chemokine release (56–58). While the temporal relationship between this glial response and neurological injury remains to be determined, available evidence suggests that astrocyte-driven inflammation may precede and shape downstream neurodegeneration (59). Therefore, the implications of acute and chronic inflammation mediated by astrocytes merit further examination in the context of alphavirus pathogenesis, particularly when assessing the risk of long-term neurological sequelae in non-lethal or subclinical infections.

This interpretation is further supported by the cytokine profile observed in brain homogenates (**Fig 11**). Infected mice exhibited a strongly pro-inflammatory milieu coinciding with peak viral replication in the CNS. Elevated levels of IFN-𝛾 observed in symptomatic mice suggest a central role in orchestrating this response, as it is known to disrupt barrier integrity, recruit immune effector cells, and alter non-immune cell function across tissues, including the CNS (60). Similarly, IL-6 has been implicated in sustained neuroinflammation and blood-brain barrier (BBB) dysfunction during viral encephalitis, often acting synergistically with TNF and IFN-𝛾 (61–63). MCP-1/CCL2, a key chemokine for leukocyte recruitment, was among the most strongly upregulated inflammatory mediators in our analysis, suggesting a prominent role in shaping the inflammatory environment. This chemokine has been shown to mediate BBB permeability and neuroinflammatory injury in alphavirus and flavivirus infections (64). Similar patterns of viral replication, glial activation, and cytokine induction have been reported in the attenuated VEEV TC-83 model, where the inflammatory milieu has been proposed to shape not only pathology but also within-host viral evolution (65). Together, these mediators may contribute to a self-reinforcing cycle of inflammation and tissue damage.

Comparable cytokine signatures have been described in models of virulent VEEV infection, where neuropathology occurs not only in regions with detectable viral antigen but also in areas of reactive gliosis, underscoring the paradoxical role of inflammation as both antiviral and pathogenic (16). *In vitro* studies further corroborate these findings, showing that VEEV-infected astrocytes release a broad array of inflammatory mediators that can promote leukocyte infiltration while influencing neuronal viability (55). Consistently, comparative studies indicate that alphavirus infection of the CNS triggers sustained induction of chemokines such as CXCL10 and CCL2/MCP-1, which have been shown to correlate with neuropathology and long-term sequelae in animal models (66). Although we did not directly assess endothelial markers, the concurrent upregulation of MCP-1/CCL2 and widespread astrocytic activation is consistent with BBB disruption. Prior studies have demonstrated that VEEV alters expression of key endothelial adhesion molecules and extracellular matrix components in the brain. Notably, ICAM-1 knockout mice show reduced neuroinflammation and delayed symptom onset, highlighting the importance of astrocyte-endothelial-leukocyte interactions in amplifying CNS injury (67). Moreover, BBB permeability may increase during later stages of infection, compounding leukocyte recruitment and inflammatory cascades even after initial viral entry (66).

The neuropathological changes observed in our model provide additional support for this sequence of events. Amino-cupric-silver staining revealed marked neurodegeneration in the cortex and striatum at 8 dpi, but not at 2 dpi, placing neuronal damage downstream of early astrocyte activation and cytokine induction (**Fig 9**). Consistently, HE-stained brain sections, including the olfactory bulb, showed meningoencephalitis with perivascular cuffing, vacuolization, and tissue disorganization (**Fig 7-8**), supporting the olfactory bulb as a major CNS entry route for VEEV after peripheral infection (53,68–70).

Taken together, our findings illustrate how an enzootic VEEV variant -traditionally considered “avirulent”- can nevertheless trigger astroglial activation, pro-inflammatory responses, and neurodegenerative damage when it encounters a permissive immunological context in a susceptible host. This aligns with clinical evidence from subtype ID circulation in Bolivia, where enzootic strains have caused dengue-like febrile illness as well as severe neurological disease in adults (2). Experimental data further show that single amino acid substitutions in VEEV glycoproteins E1/E2 can significantly alter disease progression, affecting peripheral replication, viremia levels, and CNS invasion (71). It is worth noting that a limitation of our study is the use of a laboratory-passaged RNV strain, which may exhibit enhanced neurotropism not representative of field isolates.

In sum, our study provides the first experimental characterization of RNV pathogenesis in a susceptible mouse model, revealing that this enzootic and historically neglected member of the VEEV complex can elicit significant neuroinflammatory and degenerative outcomes. By integrating diverse parameters within a single experimental framework, this work identifies key pathological checkpoints -such as cytokine elevation and splenic collapse- that warrant further mechanistic investigation. While specific aspects of RNV biology, including the precise routes of neuroinvasion or the long-term effects of lymphoid remodeling, remain to be fully elucidated, this study establishes a necessary foundational baseline. These findings challenge long-held assumptions about the clinical insignificance of RNV and illustrate how host developmental stages can unmask the pathogenic potential of otherwise neglected viruses. In a world increasingly shaped by anthropogenic disruptions, ecological compression, and expanding host–vector interfaces, viruses such as RNV likely persist within long-standing co-evolved transmission networks.

Epidemiological risk may therefore arise not from their silent circulation *per se*, but from the disruption of these fragile equilibria and from the limited capacity to detect and interpret viral activity in rapidly changing socio-ecological contexts.

## Acknowledgments

L.A.F., Y.G., and G.D.M.M. are fellows of the Consejo Nacional de Investigaciones Científicas y Técnicas (CONICET, Argentina). A.D., M.C.S., A.G., and G.A.L. are members of the Research Career of CONICET. S.D.O. and S.R.O. are members of the Technical Support Career of CONICET. This work was supported by grants from CONICET and Universidad Católica de Córdoba (UCC). The funders had no role in study design, data collection and analysis, decision to publish, or preparation of the manuscript.

## Author contributions

**Conceptualization**: Guillermo Albrieu-Llinás.

**Data curation**: Guillermo Albrieu-Llinás, Yamila Gazzoni, Glenda D. Martin Molinero, Marianela C. Serradell.

**Formal analysis**: Guillermo Albrieu-Llinás, Luciana A. Fassola, Marianela C. Serradell, Yamila Gazzoni.

**Funding acquisition**: Guillermo Albrieu-Llinás.

**Investigation**: Luciana A. Fassola, Yamila Gazzoni, Glenda D. Martin Molinero, Alicia Degano, Soledad De Olmos, Marianela C. Serradell, María E. Rivarola.

**Methodology**: Guillermo Albrieu-Llinás, Luciana A. Fassola, Yamila Gazzoni, Soledad De Olmos, Glenda D. Martin Molinero, Alicia Degano, Sergio R. Oms.

**Project administration**: Guillermo Albrieu-Llinás, Adriana Gruppi, Marta S. Contigiani.

**Resources**: Guillermo Albrieu-Llinás, Adriana Gruppi, Alicia Degano, Marta S. Contigiani.

**Supervision**: Guillermo Albrieu-Llinás, Adriana Gruppi, Alicia Degano, Marta S. Contigiani, Sergio R. Oms.

**Validation**: Yamila Gazzoni, Soledad De Olmos, Luciana A. Fassola, Guillermo Albrieu-Llinás.

**Visualization**: Yamila Gazzoni, Luciana A. Fassola, Glenda D. Martin Molinero, María F. Triquell, Soledad de Olmos.

**Writing – original draft**: Guillermo Albrieu-Llinás, Luciana A. Fassola.

**Writing – review & editing**: Guillermo Albrieu-Llinás, Luciana A. Fassola, Yamila Gazzoni, Glenda D. Martin Molinero, Soledad De Olmos, Marianela C. Serradell, Alicia Degano, Sergio R. Oms, Marta S. Contigiani, Adriana Gruppi.

## Supporting information captions

S1 Data. Raw data underlying Fig 1. Survival data used for Kaplan–Meier analyses.

S2 Data. Raw data underlying Fig 2. Individual viremia values used for kinetic analyses.

S3 Data. Raw data underlying Fig 3 Viral titers in tissues used for organ tropism analyses.

S4 Data. Raw data underlying Fig 11. Raw cytokine concentration values obtained by CBA analysis

## Notes

### Competing Interest Statement

The authors have declared no competing interest.

